# Elevated CO_2_ alters photosynthesis, growth and susceptibility to powdery mildew of oak seedlings

**DOI:** 10.1101/2023.01.07.523094

**Authors:** Rosa Sanchez-Lucas, Carolina Mayoral, Mark Raw, Maria-Anna Mousoraki, Estrella Luna

## Abstract

Elevated CO_2_ (eCO2) is a determinant factor of climate change and is known to alter plant processes such as physiology, growth and resistance to pathogens. *Quercus robur*, a tree species integrated in most forest regeneration strategies, shows high vulnerability to powdery mildew (PM) disease at the seedling stage. PM is present in most oak forests and it is considered a bottleneck for oak woodland regeneration. Our study aims to decipher the effect of eCO2 on plant responses to PM. Oak seedlings were grown in controlled environment at ambient (aCO_2_, ~ 400 ppm) and eCO_2_ (~ 1000 ppm), and infected with *Erysiphe alphitoides*, the causal agent of oak PM. Plant growth, physiological parameters and disease progression were monitored. In addition, to evaluate the effect of eCO_2_ on induced resistance (IR), these parameters were assessed after treatments with IR elicitor β-aminobutyric acid (BABA). Our results show that eCO_2_ increases photosynthetic rates and aerial growth but in contrast reduces root length. Importantly, under eCO_2_ seedlings were more susceptible to PM. Treatments with BABA protected seedlings against PM, however, this effect was less pronounced under eCO_2_. Moreover, irrespectively of the concentration of CO_2_, BABA did not significantly change aerial growth but resulted in longer radicular systems, thus mitigating the effect of eCO_2_ in root shortening. Our results demonstrate the impact of eCO_2_ in plant physiology, growth and defence, and warrant further biomolecular studies to unravel the mechanisms by which eCO_2_ increases oak seedling susceptibility to PM.

## Introduction

Plants are continually exposed to multiple biotic and/or abiotic stresses that can negatively affect their growth, productivity and survival [1, 2] Climate change, as a conjunction of carbon dioxide (CO_2_) rise, temperature increase and frequent drought periods, alters plant physiology, adaptation and resistance to pathogens [3–5]. Due to the significance of woodland habitats for biodiversity there has been a robust effort to restore, manage and defragment existing woodlands through the use of landscape level management systems [6]. Understanding how the interaction of these factors affect plant resistance responses is crucial to determine management strategies and policies to protect natural forest environments.

Pedunculate oaks (*Quercus robur*) are one of the most economically and ecologically relevant forest tree species in Europe [7]. Oaks can have life spans exceeding 900 years and this longevity allows them to reach great sizes and to sequester large amounts of CO_2_ [8]. Due to the economic and ecologic value of oaks, efforts are focusing on the regeneration of woodlands with this tree species. However, oak populations are facing several challenges that hinder their success, such as the combination of unfavourable climatic conditions and infection by pathogens [9]. One pathogen of particular relevance is the fungus *Erysiphe alphitoides*, causal agent of oak powdery mildew (PM) [10], which mainly affects young oak seedlings. *E. alphitoides* is a biotrophic pathogen present in Europe since 1907 [11] whose infection is characterised by white and cottony hyphae covering leaf surfaces, with necrosis and leaf deformation and loss at advance stage, resulting in a reduced functional leaf area and photosynthetic efficiency [12, 13]. Specifically, *E. alphitoides* causes increased transpiration rates while decreases CO_2_ assimilation and carbohydrates translocation from infected leaves to the rest of the plant [13–16]. Usually, infections result in seedling death [17] and this is why this pathogen is considered to be a bottleneck to natural woodland regeneration [18]. Sadly, there are reports as to the effect of climate on the severity of infections of PM in oak. For instance, it has been shown that higher humidity and temperatures of around 22-25°C are beneficial to pathogen growth and wet weather in late summer are beneficial for the spreading of sexual spores [19]. There is also evidence that warmer winter temperatures increase rates of infection [7]. Therefore, weather conditions associated with climate change can have a deep impact on the severity of PM disease in oak, which warrants further studies on the effect of climate change in this pathosystem.

Current methods to control PM by tree nurseries and woodland managers depend on extensive application of chemical fungicides, which use is extremely limited due to their toxicity to human health and the environment [20]. Therefore, alternative methods to control this pathogen are needed. As described in many plant species, a plausible solution could rely on the exploitation of the plant’s immune system through the activation of Induced Resistance (IR) and priming, the latter described as a sensitisation of defence mechanisms for a faster and/or stronger response upon subsequent stresses [21]. Priming can be achieved with different stimuli, including plant chemicals such as β-amino-butyric acid (BABA) [22, 23]. BABA has been shown to prime defence responses against a wide spectrum of biotic stresses [23–25] including powdery mildews in *Cucurbita* [26]. It is currently unknown whether BABA could offer a solution to protect oaks against PM. Nevertheless, it is important to consider that in agronomic settings, the use of BABA is not established as it has been shown to trigger a phytotoxic response that manifests mainly as a plant growth reduction [27, 28]. This phytotoxic response is due to the potential binding of BABA to its plant receptor [27] and to the transient relocation of energy from growth to defence [29]. It remains unknown whether BABA leads to a growth reduction in oak seedlings. Moreover, it is not understood whether the potential benefits from BABA treatment would outweigh the costs in growth in forest settings.

Associated with climate change is also the concentration of atmospheric CO_2_, which has been on the rise since the 1960’s [30]. Elevated levels of CO_2_ (eCO_2_) can drive deep changes in plant physiology and development [31]. For instance, it has been documented that eCO_2_ results in major changes in growth at early stages of plant development, primarily driven by an increased efficiency of leaves to produce biomass [32] and increased N-use efficiency [33]. In addition, under eCO_2_, starch concentrations more than double [34]. Therefore, eCO_2_ might result in enhanced growth and productivity. Whilst this could be seen as a positive “carbon fertiliser” effect, eCO_2_ has also been associated to changes in the defensive capacity of plants that have proven contrasting. For instance, in soybean, it has been described that eCO_2_ results in the production of phytoalexins and SA, resulting in enhanced resistance to pathogens [35, 36]. However, other studies testing resistance against fungal pathogens have described that eCO_2_ does not impact resistance in ragweed [37] or even increases susceptibility in wheat [38]. Specific studies have been done to test the effect of eCO_2_ in PM diseases. eCO_2_ results in increased disease severity in Arabidopsis [39], soybean [40] or grapevine [41]. However, no effects were found in barley [42], or zucchini [43]. Interestingly, a study in Arabidopsis has reported that changes in the CO_2_ concentration alter the capacity of plants to express IR [5]. The effect that eCO_2_ has in the severity of PM and expression of IR in oak remains to be studied.

Despite the extensive knowledge on the energy relocation trade-off between plant growth into defence [44] [45], little is known whether an increase in growth can also divert energy away from the activation of resistance responses. Nevertheless, it could be hypothesised that eCO_2_ would increase growth through the production of young and tendered leaves, which could be more susceptible to the fungus [12], thus resulting in a detrimental effect. In this study, a range of physiological parameters and disease phenotypes have been linked to changes in growth triggered by eCO_2_ and BABA, thus assessing the impact of eCO_2_ on oak PM infections and IR.

## Materials and methods

### Plant material and growth conditions

Acorns of *Quercus robur* 403 UK provenance [46] were germinated according to existing protocols [47, 48]. After 72 h, germinated acorns were transferred into individual root trainers (Maxi Rootrainers, Haxnicks, RT230101) containing 400 mL of Scott’s M3 soil. Germinated acorns were transferred into the growth chambers (CONVIRON, Controlled Environments, Inc., Winnipeg, Canada) and grown at 16/8 h light day/night 20°C/17°C cycle and irrigated at field capacity throughout the experiment. Two different CO_2_ concentrations were used: ~400ppm for ambient CO_2_ (aCO_2_) and ~1000ppm for elevated CO_2_ (eCO_2_). After initial experiments, germination of plants exposed to eCO_2_ was delayed 7 days with respect to the aCO_2_ plants to allow for similar developmental stages upon infection.

### B-aminobutyric acid treatment

β-amino-butyric-acid (BABA) was purchased from Sigma (Catalogue number A44207). Treatment solutions were freshly prepared on the day of treatment. BABA treatments were performed entirely as previously described for other plant species [25]. Four-week-old seedlings were soil-drenched with 40 mL per plant of a 50 mM BABA solution, resulting in a final concentration in the soil of 5 mM. Control treatment plants were treated with water. After treatment, watering was done in equal amounts to maintain BABA concentrations and irrigation status stable among plants and treatments.

### Pathogen infection and disease scoring

Powdery mildew (PM) causal agent *E. alphitoides* was cultivated and maintained on oak seedlings. Spores were collected by shaking infected leaves in water and 0.05% Silwet-L77 (CAS 27306-78-1, De-Sangrosse). Inoculations were performed by spraying inoculum containing between 1.5 and 1.7 × 10^6^ spores/mL onto leaves of the entire oak seedlings. After inoculation, plants were covered with plastic bags to maintain high relative humidity and placed back into the growth chambers. Non-inoculated controls were sprayed with a solution of water and 0.05% Silwet-L77. Disease was scored by counting the number of colonies per leaf, the number of leaves affected (i.e. presenting colonies) and the number of discarded leaves (“lost”), the latter parameter assessed by counting the number of leaf nodes without leaves.

### Leaf physiological measurements

Physiological measurements were conducted with a portable open gas exchange system (LI-6400XT, LI-COR, Lincoln, NE, USA) at three campaigns (time points) representing different infection stages: Campaign 1 at 0 days post infection (dpi), before infection - no symptoms; Campaign 2 at 7 dpi; and Campaign 3 at 14 dpi. Measurements were done consistently on the same leaf, which had been produced on the first leaf flush. Photosynthetic rate (An) and stomatal conductance (gs) were measured at a saturating light intensity of 700 μmol photons m^-2^s^-1^ (as determined by light response curves of An) and a flow rate of 300 μmol s-1. Reference CO_2_ was set at either 1000 μmol mol□^1^ eCO_2_ or 400 μmol mol-^1^ for seedlings grown under elevated or ambient CO_2_, respectively. Maximum photochemical efficiency of photosystem II (Fv/Fm) was measured at the final experimental point (14 dpi) using a portable Pocket PEA chlorophyll fluorometer (Hansatech Instruments Ltd) in the same leaves used for gas exchange measurements after a 30-minutes dark-adaptation period [49].

### Aerial plant growth parameters

Height and main stem diameter were used to assess growth. Height was determined with a ruler (30cm, Helix L16) by measuring the length from the ground to the top of the main stem. Diameter was determined with a calliper (0.01 mm; Carbon Fiber Composite Digital Calliper, EAN 5421815474575) at a specific and consistent point of the main stem at the second node after cotyledons. Relative growth rates (RGRs) in height (HRGR) and main stem diameter (DRGR) were determined between Campaigns 1 and 2 and Campaigns 2 and 3, corresponding with 0 and 7 dpi and 7 and 14 dpi, respectively, using the formula: RGR= (lnXt_2_-lnXt_1_)/(t_2_-t_1_), where Xt_2_ and Xt_1_ represent seedling height (cm) or shoot diameter (mm) at time points t_2_ and t_1_ [50].

### Root plant growth parameters

Plant material was collected at Campaign 3 (14 dpi) to perform destructive measurements. Root length was measured as the distance between the hypocotyl and tip of the primary root using a ruler (30cm, Helix L16).

### Biomass parameters

Biomass was calculated with the fresh and dry weight of roots and shoots at Campaign 3 (14 dpi). Material was collected in aluminium foil, weighted (fresh weight, FW), dried for two days in a 60°C oven until constant mass and weighed (dry weight, DW) with a laboratory scale (KERN EMB600-2).

### Statistical analyses

Statistical and data analyses were done using PROC MIXED (SAS 9.4 Institute, Inc.). Eight plants (four per CO_2_ condition) were used for non-inoculated controls and this treatment was named “Non-inoc”. Sixteen plants (eight per CO_2_ condition) were used for the inoculated group treated with water and these were named “Inoc-Water”. Similarly, sixteen plants (eight per CO_2_ condition) were used for the inoculated group treated with BABA and this treatment was named “Inoc-BABA”. Analysis of Variance was performed to test differences among treatments using threshold p<0.05. Tukey post-hoc test at p<0.05 was used to compare means. In all analyses, residual plots were generated to identify outliers and to confirm that variances were homogeneous and normally distributed. Experiments on disease assessment and growth have been repeated twice with similar results.

## Results

### Effect of elevated CO_2_ on growth and disease resistance

Seedlings grown under eCO_2_ were taller than the ones growing at aCO_2_ until 2 weeks of growth when heights stabilised (Fig. 1A; Fig. S1A). Experiments revealed that plants growing under eCO_2_ presented a higher number of PM colonies (Fig. 1B; Fig. S1B) and a higher number of leaves affected by the disease (Fig. 1C). Therefore, eCO_2_ results in enhanced initial plant growth and susceptibility to PM.

**Figure 1.**
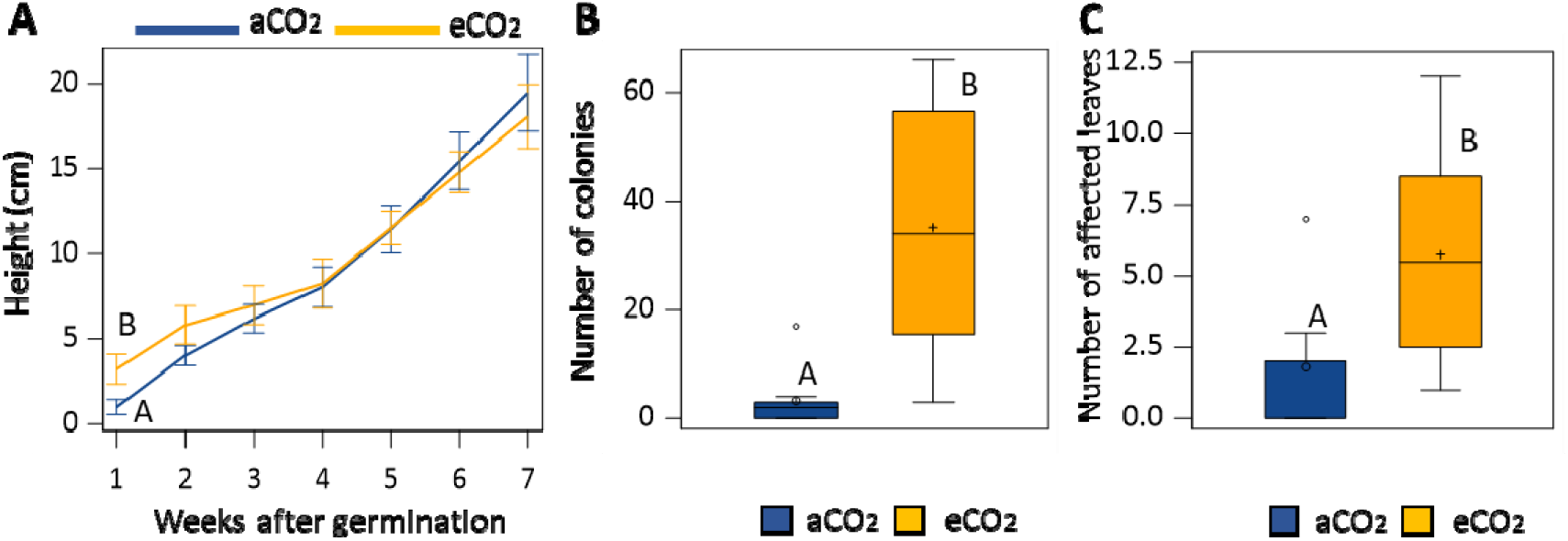
Effects of eCO_2_ on growth **(A)** and PM resistance represented by the number of colonies per plant **(B)** and the number of leaves affected (with symptoms i.e. mycelium and/or necrosis) **(C)** at 14 days post infection (dpi). Capital letters represent statistically significant differences between CO_2_ concentrations (Tukey post-hoc test; p<0.05; n=8 for inoculated plants/n=4 for non-inoculated plants).

### Effect of B-amino butyric acid (BABA) on growth and induced disease resistance

Seedlings treated with BABA presented lower height values than plants treated with water (Fig. 2A), however this observation was not statistically significant (p-values ranged from 1 to 0.052). Treatments with BABA resulted in fewer colonies (Fig. 2B) and a reduced number of affected leaves (Fig. 2C) than water-treated plants. Therefore, BABA enhances resistance against PM while potentially reducing plant growth.

**Figure 2.**
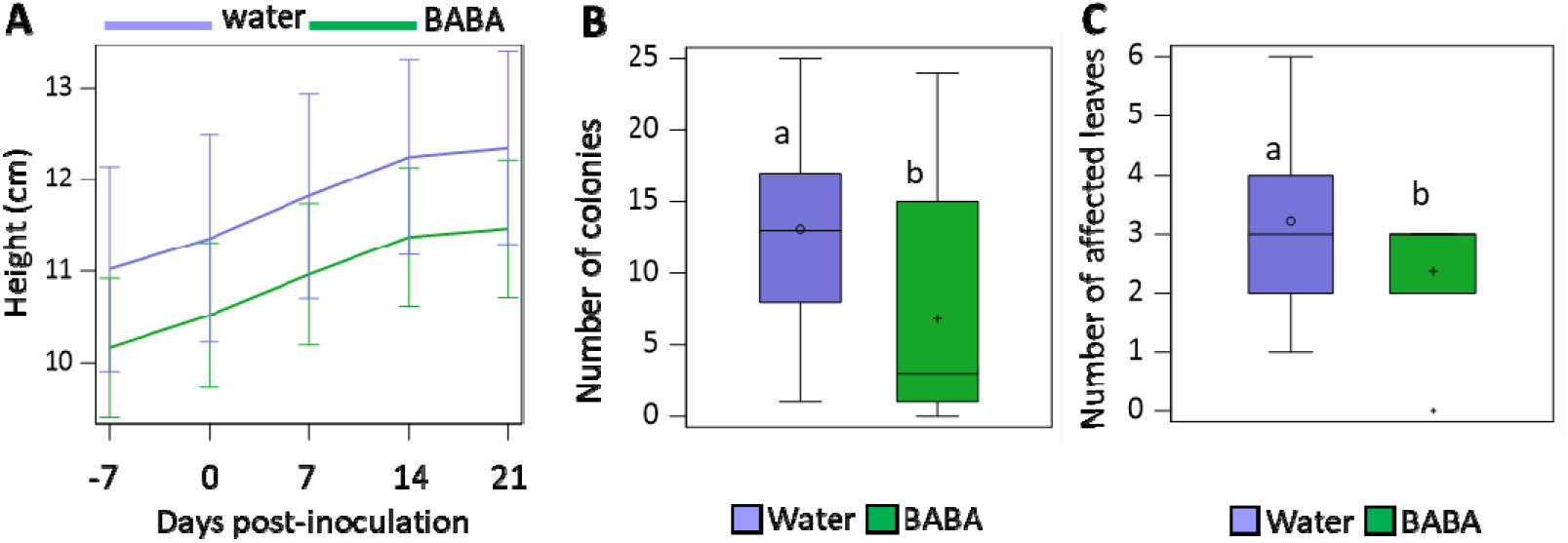
Effects of BABA treatment on growth **(A)** growth **(A)** and PM resistance represented by the number of colonies per plant **(B)** and the number of leaves affected (with symptoms i.e. mycelium and/or necrosis) **(C)** at 14 days post infection (dpi). Lowercase letters represent statistically significant differences between BABA and water treatments (Tukey post-hoc test; p<0.05; n=8 for inoculated plants/n=4 for non-inoculated plants).

### Effect of eCO_2_ on BABA-induced resistance against powdery mildew

To assess the impact of eCO_2_ on BABA-induced resistance (IR) against PM infection, plants growing at aCO_2_ and eCO_2_ were treated with water or BABA one week before infection with PM. To have plants of similar developmental stages upon treatment and infection, eCO2 plants were germinated a week later [5]. As seen before, under aCO_2_, BABA induces resistance against PM as BABA-treated plants have a lower number of colonies and leaves affected (Fig. 3A and B). eCO_2_ treatment in this particular experiment, resulted in a dramatic enhanced susceptibility to PM that manifested as a high number of leaves being discarded by the plant (Fig. 3C - “lost” leaves). Consequently, the number of colonies and affected leaves in infected plants under eCO_2_ was lower than the aCO_2_ plants (Fig. 3A and B). Water-treated seedlings discarded leaves under both CO_2_ conditions and BABA treatments fully protected from severe disease with no discarded leaves observed (Fig. 3C). Interestingly, eCO_2_-grown plants treated with BABA had a higher number of colonies and affected leaves than aCO_2_-grown BABA treated plants, however these observations were not statistically significant. Therefore, BABA-induced resistance is maintained under eCO_2_, but our results suggest that the treatment may not be as effective under eCO_2_ as it is under aCO_2_.

**Figure 3.**
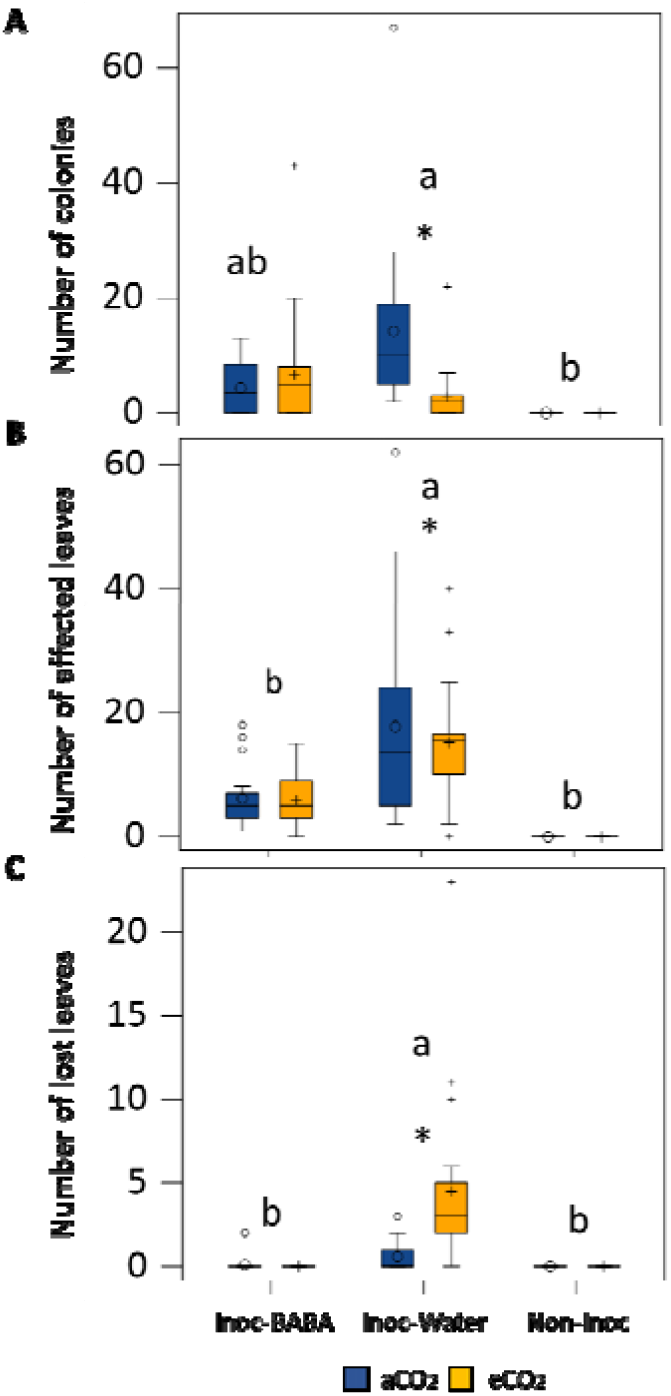
Combined effect of eCO_2_ and BABA on PM disease. Graphs indicate number of colonies **(A)**, number of affected leaves due to the infection **(B)** and number of lost leaves at the end of the experiment (14dpi) **(C)**. Lowercase letters inside the figures represent statistically significant differences between groups: Inoc-BABA representing infected seedlings treated with BABA; Inoc-water representing infected seedlings and Non-inoc representing non-infected seedlings. Asterisks indicate statistical differences between CO_2_ concentration (Tukey post-hoc test; p<0.05; n=8 for inoculated plants/n=4 for non-inoculated plants).

### Effects of eCO_2_ and BABA treatment on physiological parameters

We found that eCO_2_ overall significantly increased An of infected seedlings throughout the experimental period (Fig. 4A) compared to aCO_2_. However, non-inoculated seedlings did not show differences in An between CO_2_ levels in Campaigns 2 and 3. Under eCO_2_, differences between BABA/water treatment and inoculated/non-inoculated seedlings were not detected (Fig. 4A). However, under aCO_2_, significant differences were found between BABA and water-treated seedlings as well as with the non-inoculated seedlings: BABA-treated seedlings maintained higher An than water-treated seedlings and in Campaign 3, non-inoculated seedlings had the highest An (Fig. 4A).

**Figure 4.**
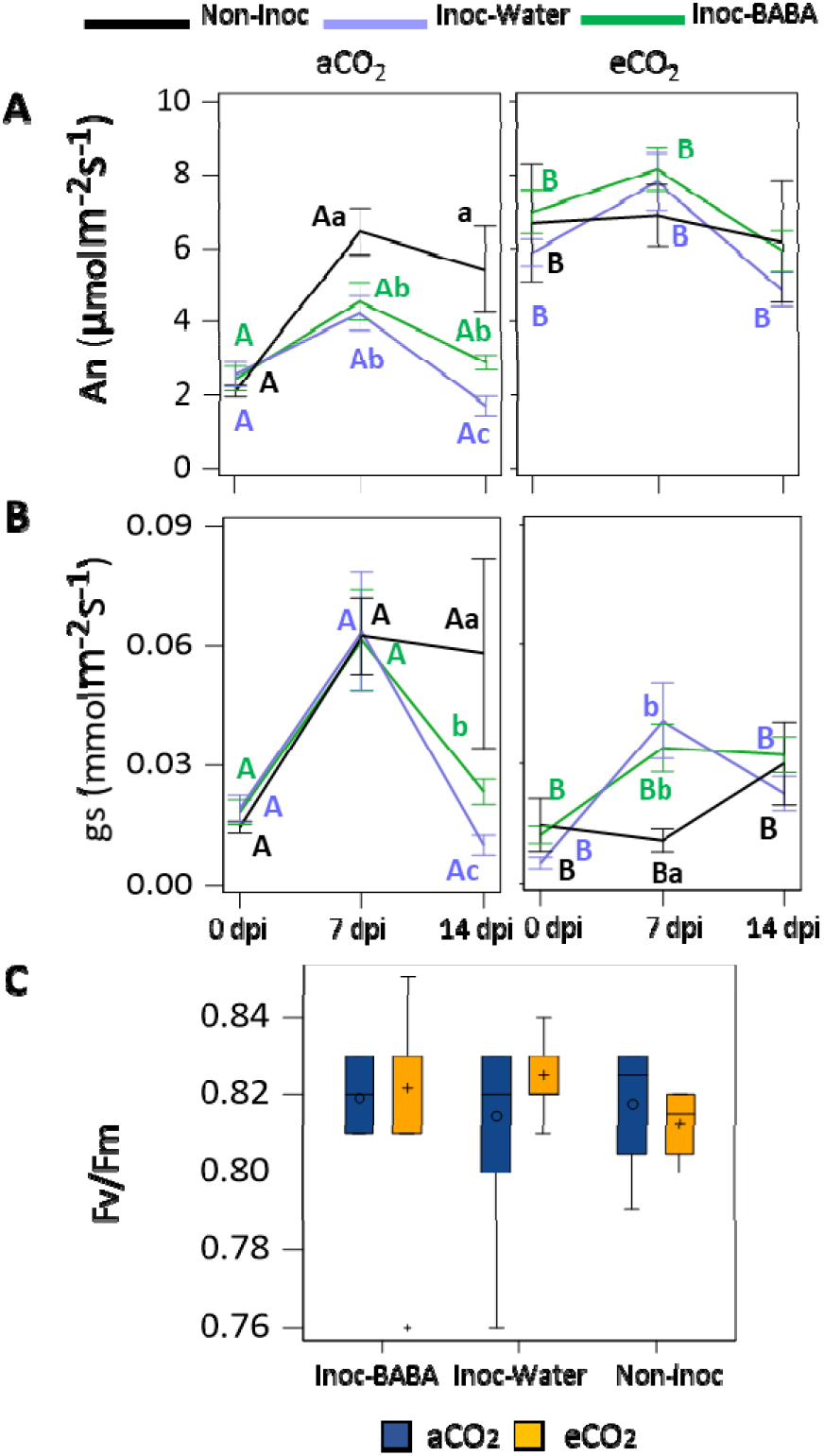
Combined effect of elevated CO_2_ levels and BABA treatment on physiological parameters. Net photosynthetic rates (An) **(A)**, Stomatal conductance (gs) **(B)** and Photosynthetic Efficiency of Photosystem II **(C)**. The horizontal axis represents the measurement campaigns at 0, 7 and 14 dpi. Lowercase letters inside the figures represent statistically significant differences between groups: Inoc-BABA (green) representing infected seedlings treated with BABA; Inoc-water (light blue) representing infected seedlings and Non-inoc (black) representing non-infected seedlings. Capital letters indicate statistical differences between CO_2_ concentration (Tukey post-hoc test; p<0.05; n=8 for inoculated plants/n=4 for non-inoculated plants).

Stomatal conductance (gs) was measured in all treatments. In contrast with the results observed for the An, seedlings growing under eCO_2_ had a trend of lower values of stomatal conductance (Fig. 4B). This observation was more pronounced in non-inoculated seedlings in Campaigns 2 and 3. Under eCO_2_, no statistically significant differences were found between treatments. Under aCO_2_ conditions, similar profiles to those for An were observed, with noninoculated plants showing the higher stomatal conductance, followed by inoculated plants treated with BABA and then inoculated plants treated with water (Fig. 4B).

Minor differences were found for maximum photochemical efficiency of photosystem II (Fv/Fm) with the different concentrations of CO_2_: plants growing under eCO_2_ had slightly lower levels of Fv/Fm (Fig. 4C). However, no significant differences were found at either treatment or infection profile (Fig 4C).

### Effects of eCO_2_ and BABA treatment on growth

Height, main stem diameter, root length and root and shoot biomass were measured to test the effect of eCO_2_ and BABA on growth. eCO_2_-grown plants were germinated one week after aCO_2_-grown plants to work with plants at similar developmental stages guided by results presented in Fig 1A. Accordingly, no differences in height were observed between aCO_2_ and eCO_2_ plants (Fig. 5A). However, height RGR (HRGR) revealed differences throughout the experiment. Firstly, between Campaigns 1 and 2, both water and BABA-inoculated plants showed higher HRGRs under eCO_2_ in comparison to their respective controls growing under aCO_2_. Second, this effect was more pronounced in water treated plants under eCO_2_ as they had a higher HRGR which was statistically significant compared to the HRGRs of BABA and non-inoculated plants. Third, as the experiment progressed, HRGRs decreased in all treatments apart from in plants treated with BABA and grown under aCO_2_ (Fig. 5B). Similarly to the height measurements, there were not statistically significant differences between aCO_2_ and eCO_2_ diameters (Fig. 5C). Within CO_2_ concentrations, BABA-treated plants inoculated with PM had bigger diameters than the water-treated plants and noninoculated plants at Campaigns 1 and 2. At Campaign 3 no differences in diameters were found between treatments. RGR values of the diameter (DRGR) showed that those observed under eCO_2_ were higher between Campaigns 2 and 3 in non-inoculated plants and in water-treated inoculated plants (Fig. 5D), however no differences were found in BABA-treated plants. Interestingly, whereas DRGR values were reduced in all treatments as the experiment progressed, the values of non-inoculated plants increased under eCO_2_. (Fig 5D).

**Figure 5.**
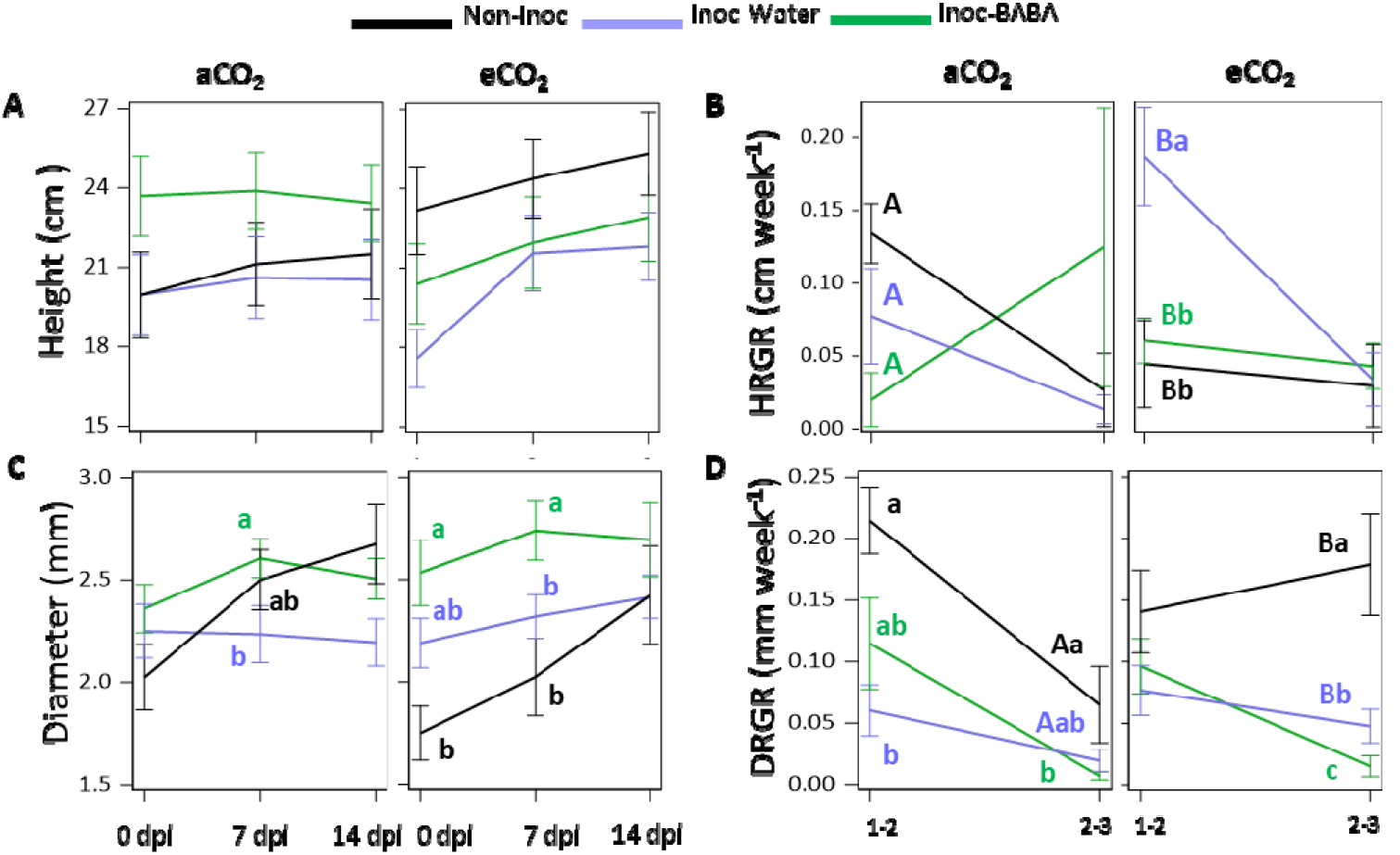
Combined effect of eCO_2_ levels and BABA treatment on morphological parameters. Plant height **(A)**, height relative growth rate (HRGR) **(B)**, trunk diameter **(C)** and trunk relative growth rate (DRGR) **(D)**. The horizontal axis represents the measurement campaigns at 0, 7 and 14 dpi. Lowercase letters inside the figures represent statistically significant differences between groups: Inoc-BABA (green) representing infected seedlings treated with BABA; Inoc-water (light blue) representing infected seedlings and Non-inoc (black) representing non-infected seedlings. Capital letters indicate statistical differences between CO_2_ concentration (Tukey post-hoc test; p<0.05; n=8 for inoculated plants/n=4 for non-inoculated plants).

At the final point right after measuring Campaign 3, destructive measurements in root length and root and shoot biomass were recorded. BABA-inoculated seedlings presented higher root length than non-inoculated and water inoculated seedlings (Fig. 6A) and this observation was more pronounced under aCO_2_. eCO_2_ resulted in shorter roots in both inoculated water- and BABA-treated seedlings (Fig. 6A) however no differences were found in non-inoculated seedlings. Similar patterns were observed in root biomass and in root length as BABA-treated seedlings had heavier roots systems (Fig 6B). Interestingly, no significant differences in root biomass were found between CO_2_ concentrations. Shoot biomass revealed no statistically significant differences between aCO_2_ and eCO_2_ or between treatments (Fig. 6C).

**Figure 6.**
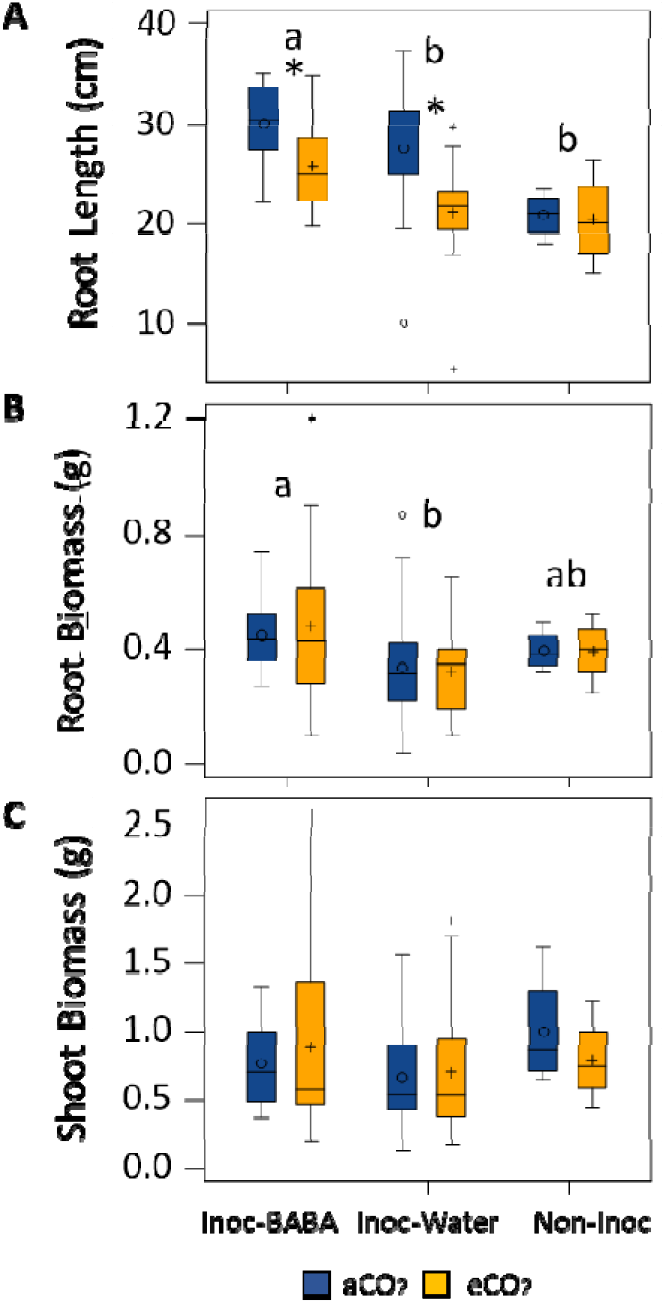
Combined effect of eCO_2_ levels and BABA treatment on root traits. Graphs represent root length **(A)**, root biomass **(B)** and shoot biomass **(C)** at the end of the experiment (14 dpi). Lowercase letters inside the figures represent statistically significant differences between groups: Inoc-BABA (green) representing infected seedlings treated with BABA; Inoc-water (light blue) representing infected seedlings and Non-inoc (black) representing non-infected seedlings. Asterisks indicate statistical differences between CO_2_ concentration (Tukey post-hoc test; p<0.05; n=8 for inoculated plants/n=4 for non-inoculated plants).

## Discussion

We have attempted to disentangle growth and defence phenotypes caused by elevated CO_2_ (eCO_2_). Even when it could be considered as a plant growth stimulant (e.g. increased stem growth, woody biomass, leaf area index and root number), eCO_2_ has been demonstrated to trigger a stress response in plants [51, 52] and shown to affect the plant’s defence capacity, although in the latter, highly contrasting phenotypes have been observed [39, 41, 53]. We have identified that worryingly, eCO_2_ results in enhanced susceptibility to the already damaging pathogen causing PM disease (Fig. 1B, 1C, 1D, Fig. 3 and Fig. S2). This is in line with the results of Lake and Wade [39] and Khan and Rizvi [54] as they observed the increased susceptibility to cucurbit PM *Erysiphe cichoracearum* under eCO_2_ (800 ppm and ~600 ppm, respectively). This increased susceptibility was associated with a higher colonisation, changes in stomatal abundance and foliar thickness. In agreement with this, other researchers have also shown enhanced susceptibility to PMs after exposure to eCO_2_, however, they demonstrated that the susceptibility phenotype only occurred when CO_2_-exposed plants were also grown under high temperatures [41, 43]. Considering that temperatures in the planet are expected to increase considerably, and that temperature is a determinant factor in the virulence of PM-causing pathogens [7] our results on the effect of eCO_2_ in increased susceptibility to PMs warrant attention. Whereas high temperatures and eCO_2_ trigger even higher susceptibility to PM in oak seedlings remains unknown. However, considering our results and previous literature, it is plausible to speculate that the added stress of high temperatures could result in an even further pronounced susceptibility phenotype. Mechanistic explanations linking eCO_2_ and enhanced susceptibility are based on the observations that eCO_2_ alters the plant concentrations of salicylic acid (SA), jasmonic acid (JA) and/or polyphenols [5, 54–56]. All these compounds are directly involved in plant defence mechanisms [57, 58]. However, the effects of eCO_2_ in leaves have been shown to be more complex: plant resistance mechanisms are more than changes on levels of secondary metabolites and hormones [59, 60]. For example, Teng et al. [61] described reduced stomatal density and stomatal index of leaves under 700 ppm associated with susceptible phenotype. In contrast, in Arabidopsis, eCO_2_ has been shown to affect cell-wall defence, through the enhanced deposition of callose upon insect attack [62, 63], the up-regulation of a key gene involved in callose biosynthesis [64] and the increased production of starch [34], a key regulator of callose [65, 66]. Therefore, these results in Arabidopsis suggest that eCO_2_ triggers enhance callose deposition, which is correlated with a phenotype of enhanced resistance. Considering that there is evidence of an effect of eCO_2_ in cell-wall defence, we can speculate that in oak seedlings, exposure to eCO_2_ could reduce callose and consequently result in enhanced susceptibility.

The exploitation of the plant immune system through induced resistance (IR) has been proposed as an alternative for plant disease control. The IR compound β-amino butyric acid (BABA) provides protection in a variety of plants against PM diseases [23–26]. BABA enables defences by priming callose deposition and the activation of the SA-dependent defence pathways [24, 25, 28, 67]. The mechanisms by which BABA-IR manifests in oak seedlings against this biotrophic pathogen are currently under study, however we are able to hypothesise that as described in other species, BABA will induce resistance through priming of SA and callose [68], which are known to be highly effective against PMs [69, 70]. Our results indicate that BABA under eCO_2_ performs worse than under aCO_2_, although this phenotype was not statistically significant (Fig. 3). However, this observation could serve us to hypothesise different mechanisms of action of BABA-IR in oak seedlings. Considering that eCO_2_ has been shown to increase levels of SA and that eCO_2_ primes SA-dependent gene expression (PR1) [5], it is easy to speculate that the potential reduced effect of BABA under eCO_2_ would be determined by other BABA-IR mechanisms such as cell-wall defence by altering priming of callose deposition. However, considering the evidence on priming of callose deposition by eCO_2_ in Arabidopsis [62, 63], these observations represent a potential contrasting phenotype to the reduced effect of BABA in oak seedlings, thus pointing towards the effect of eCO_2_ on priming to be plant species-specific. Further investigations are therefore needed to unravel the effect of eCO_2_ in IR and priming of oak seedlings.

Enhanced photosynthesis by eCO_2_ has been described in multiple studies [31, 51, 71]. We found that eCO_2_ increases photosynthesis rates at the same time that reduces stomatal conductance, two key physiological parameters that influence plant growth, development and biomass production [71]. According to our results, in aCO_2_ conditions, stomata were strongly closed at 0 dpi and that is why the conductivity and An are so low. Plants grown at eCO_2_ show an even lower conductivity while also showing higher An, which could be because CO_2_ concentration in the chloroplast and intercellular spaces was very high, cells were saturated with CO_2_, and could therefore keep their stomata more closed and photosynthesise more than plants grown under aCO_2_. This consequently would also result in a higher efficiency of water use, as we could see in our experiment with the extremely high values of An and low values of gs (Fig. 4A and 4B). In addition, we did not observe statistically significant changes in photosynthetic efficiency of PSII (Fv/Fm) between treatments, although eCO_2_ seemed to reduce this parameter slightly, which could be associated to the known induced stress of eCO_2_ [72–74]. Moreover, in general, increased concentrations of CO_2_ trigger enhanced growth in plants [53]. Indeed, we have observed increases in HRGR and DRGR triggered by eCO_2_ (Fig. 5B and 5D).

Some studies have focussed on evaluating changes in physiological and growth parameters upon biotic attacks under eCO_2_. For instance, infection with PM of barley plants showed that RGRs of shoot and root dry weight were higher in PM infected plants growing at eCO_2_ in comparison to plants grown at aCO_2_, attributing that faster growth to the increased An [42]. Similarly, we found higher HRGR growing under eCO_2_ at early stages of infection and higher DRGR at later stages of infection (Fig. 5B and 5D). This correlated with higher levels of An under eCO_2_ throughout the experiment (Fig. 4A). The enhanced growth patterns at different stages of infection allow us to speculate that eCO_2_ accelerates developmental processes as, naturally, seedlings invest first in aerial growth towards leaf production and then the engrossment of trunks. Importantly, however, only in aCO_2_ grown plants we did observe a reduction in An and stomatal conductance after infection. Moreover, the increase in DRGR was much more pronounced in non-inoculated plants growing under eCO_2_ than in inoculated plants. This is likely due to (i) a reduced net assimilation rate in PM-infected plants through a source effect, i.e. PM is directly responsible for a decrease in photosynthesis like documented in other foliar diseases [75] or; (ii) to the disease itself, caused by an obligate biotroph, which can also reduce the net assimilation rate through a sink effect, i.e. PM diverts carbon fluxes from growing plant organs to themselves for their own growth [76, 77]. Nevertheless, no changes were observed in An and stomatal conductance in inoculated seedlings growing under eCO_2_. Thus, eCO_2_ exposure at the levels used in our experiments results in enhanced physiological parameters irrespectively of the infection status.

BABA-IR is known to be coupled to a growth reduction in herbaceous plants due to the binding of BABA to its plant receptor [27] or/and the relocation of resources from growth to defence [29]. Our results confirm that BABA enhances resistance in oak seedlings against PM with a slight detriment in growth (Fig. 2A; Fig. 5B). In contrast, BABA treatment had no significant impact on physiological parameters (Fig. 4A, 4B) at early infection stages. This suggests that the impact of BABA in growth does not occur while the infection has not fully developed. Interestingly, however, we observed that after infection, plants treated with BABA had physiological and growth values resembling closer to the ones of non-inoculated plants than the water infected ones in almost all parameters (Fig. 3 to Fig. 6). We consider this to be a buffering effect of BABA on the impact of the infection in physiology. Importantly, this buffering effect was again more pronounced in plants grown under aCO_2_ than the ones grown under eCO_2_, thus showing the clear driver effect of eCO_2_ in physiology and providing a potential explanation of the reduced effect of BABA-IR under eCO_2_. A reason for why this is happening could be associated with an increased C:N ratio of plant biomass caused by exposure to eCO_2_ [52, 78, 79]. Fertirrigated basil plants with NH4+ have been shown to have less severe downy mildew disease [80]. Importantly, studies on *Vicia* and *Medicago* have shown that BABA treatments result in an increased N levels in leaves [81]. Therefore, it is easy to hypothesise that BABA reduces C:N ratios in oak seedlings to trigger effects on resistance to foliar diseases, which could explain part of the BABA-protective role as well as the weaker effect of BABA under eCO_2_.

At the end of the experiment, root and shoot traits were assessed and compared. We found no differences in shoot biomass under eCO_2_ in any of the treatments, which could be because mycelium or cumulative spore biomass can represent up to 50% of infected leaf biomass [76]. Similar lack of differences on dry mass were also observed on soybean plants infected with PM and grown under eCO_2_ [40]. In addition, no differences were found in root biomass between aCO_2_ and eCO_2_. Interestingly, however, we observed a significant reduction of root length under eCO_2_ with respect to aCO_2_-grown seedlings after infection. Therefore, in the presence of an infection under eCO_2_, roots were shorter but they had similar dry weights. To explain this observation, we need to revise the process of carbon (C) allocation to maximise trade-offs between growth and defence [82]. CO_2_ fixed by photosynthesis in chloroplasts has several possible fates, but a considerable part of it ends up as sucrose, which is consumed in respiration and growth or is stored as solutes in vacuoles. Sucrose excess as result of eCO_2_ can be exudated through the roots as organic acid derivatives such as malic acid [83], which in turn could acidify the soil surrounding the plant under eCO_2_ and damage the tissue thus resulting in shorter roots. Surprisingly, soil-drench BABA treatment produced longer and heaviest roots than the water-treated controls (Fig 6). This result could be due to the biostimulant role of amino acids for yield and growth improvements [84–86]. This biostimulatory effect of amino acids is caused by the modulation of plant molecular and physiological processes [87], including direct effects on C and nitrogen (N) metabolism and N uptake [88–90]. However, even when eCO_2_ also triggered a reduction of the root length in BABA-treated seedlings upon infection, this phenotype was less pronounced than in inoculated water-treated plants. Considering the demonstrated effects of eCO_2_ on the levels of defence compounds (e.g. SA) [5, 53–56], the reduced effect of BABA under eCO_2_ against PM and the mechanisms of action of BABA in other plant species (e.g. priming of SA-dependent defences) [25, 67, 91, 92], we could speculate that there are trade-offs between growth and defence driven by eCO_2_.

The molecular mechanisms underlying growth and defence trade-offs under eCO_2_ remain to be elucidated. Nevertheless, our results have shown that whereas eCO_2_ increases photosynthesis and growth, it also enhances susceptibility to PM and partially hinders the effect of BABA, a plausible method of control of this disease in tree nurseries. Therefore, considering that oak trees are keystone trees of our European forests and that their regeneration is threatened by the high susceptibility of oak seedlings to PM disease, our results warrant further investigations on the risks of Climate change associated with enhanced levels of atmospheric CO_2_.

## Acknowledgments

This work has been supported by the JABBS foundation project “Resistance strategies of oak trees in the arms race with pathogens” to EL. We are extremely grateful to the Birmingham Institute of Forest Research (BIFoR) for providing the equipment required for the physiological parameters reported here.

## Data Availability Statement

All supporting data are included within the main article and its supplementary files.

## Author contribution

EL conceived and designed the study and obtained core funding. RSL and CM design the experimental work pipeline. RSL, CM, MR and MAM, conducted experiments and gathered data. RSL and CM designed the data analysis pipeline. CM performed statistical analyses and data interpretation with assistance from RSL. RSL, CM and EL wrote the article with input from all authors.

