## Supplementary material for "Elevated CO_2_ alters photosynthesis, growth and susceptibility to powdery mildew of oak seedlings": Fig S1

Supplem**entary Figure S1**. Visual effects of enhanced CO2 on A) growth at early timepoints and B) powdery mildew resistance in oak represented

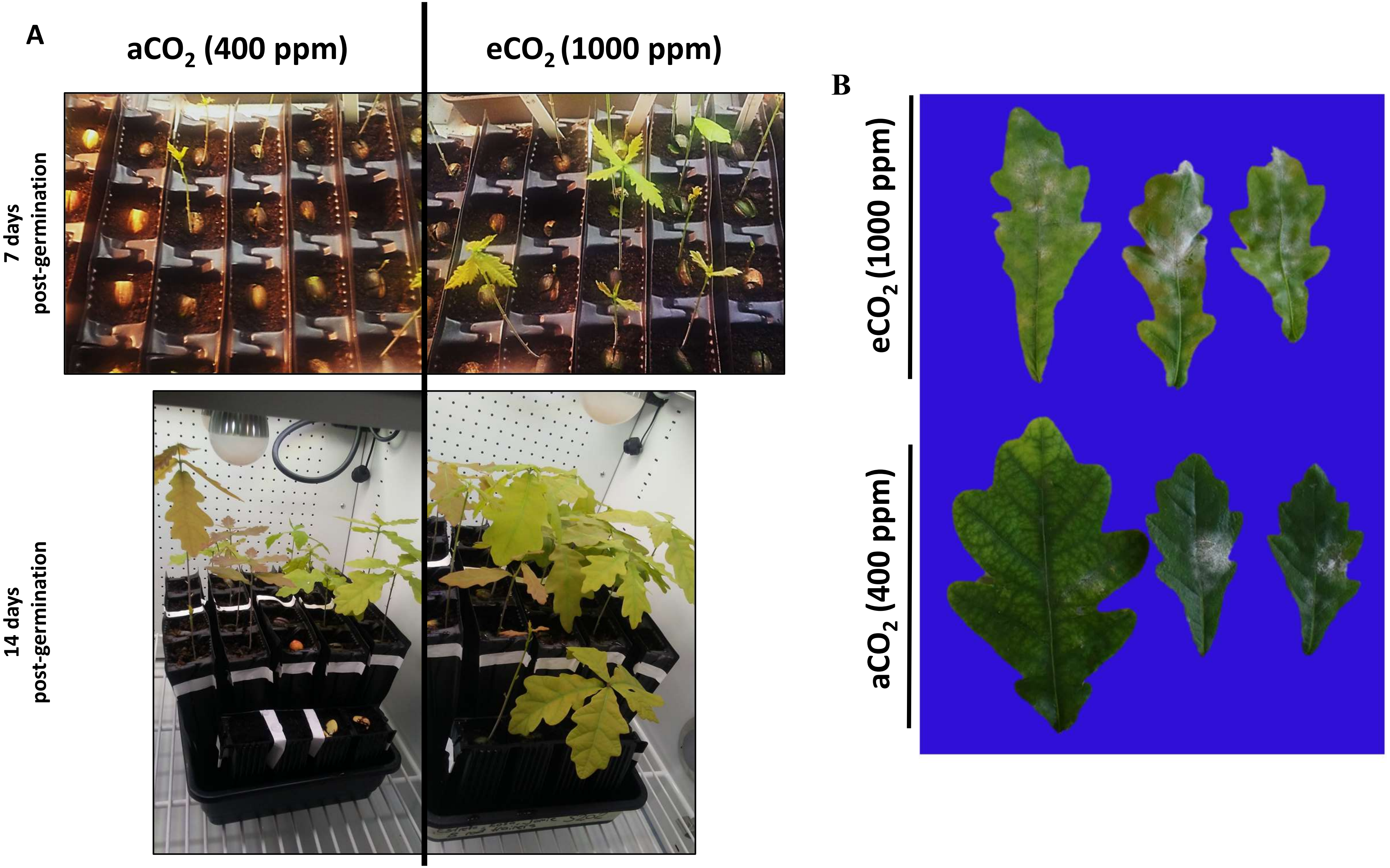
