## Supplementary material for "Elevated CO_2_ alters photosynthesis, growth and susceptibility to powdery mildew of oak seedlings": Fig S2

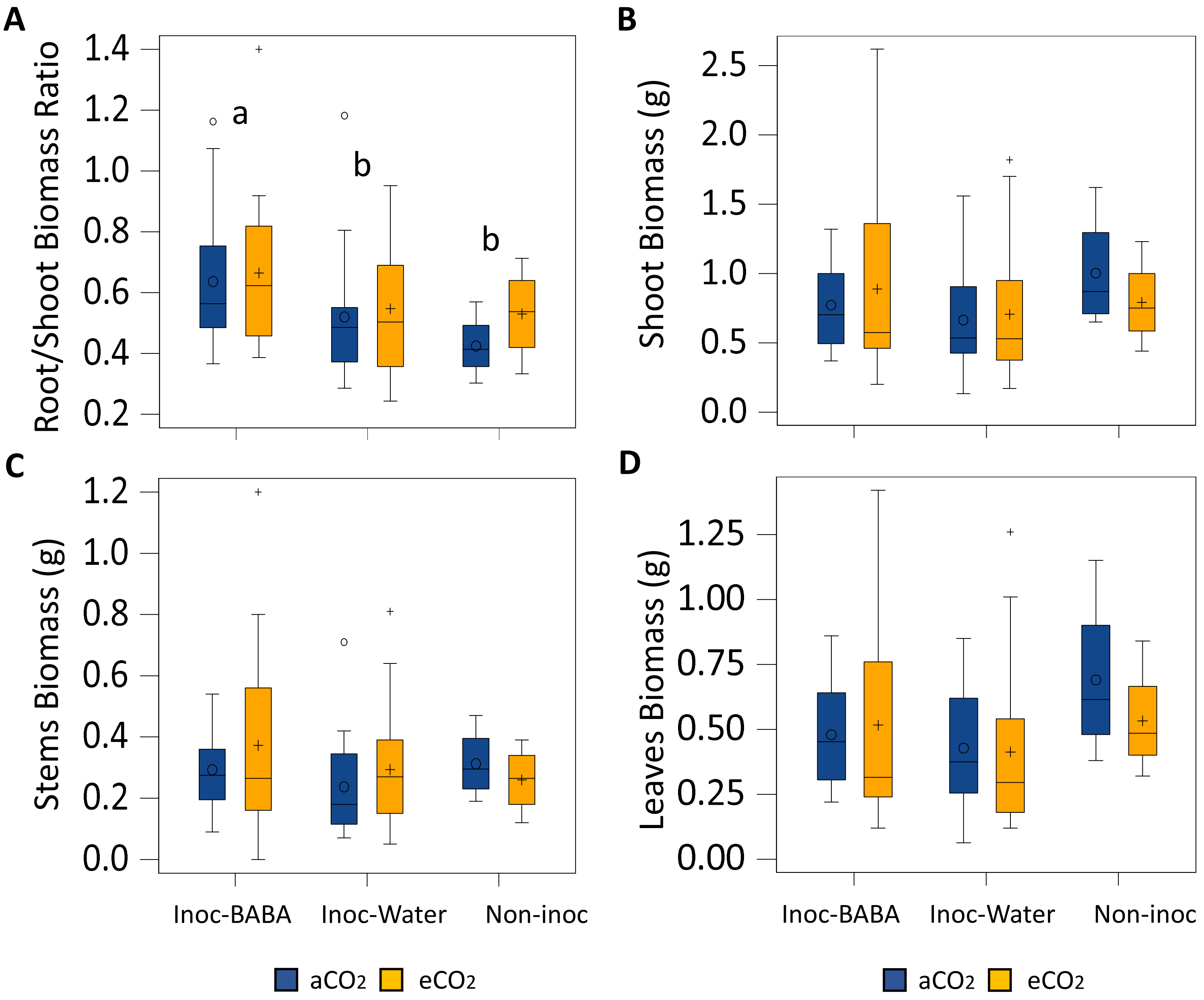


**Supplementary Figure S2.** Combined effect of enhanced CO2 levels and BABA treatment on biomass allocation in roots/shoot ratio **(A)**, shoots **(B)**, stems **(C)** and leaves **(D)** at the end of the experiment. Lowercase letters inside the figures represent statistically significant differences between groups: Inoc-BABA representing infected seedlings treated with BABA; Inoc-water representing infected seedlings and Non-inoc representing non-infected seedlings (Tukey post-hoc test; $p<0.05$; $n=8$ for inoculated plants/$n=4$ for non-inoculated plants).
