## Supplementary material for "Elevated CO_2_ alters photosynthesis, growth and susceptibility to powdery mildew of oak seedlings": Fig S3

**Supplementary Figure S3**. Combined effect of enhanced CO2 and BABA treatment on oak seedlings phenotypes. Inoc-BABA represents infected seedlings treated with BABA; Inoc-Water represents infected seedlings treated with water and Non-inoc represents non-infected seedlings.

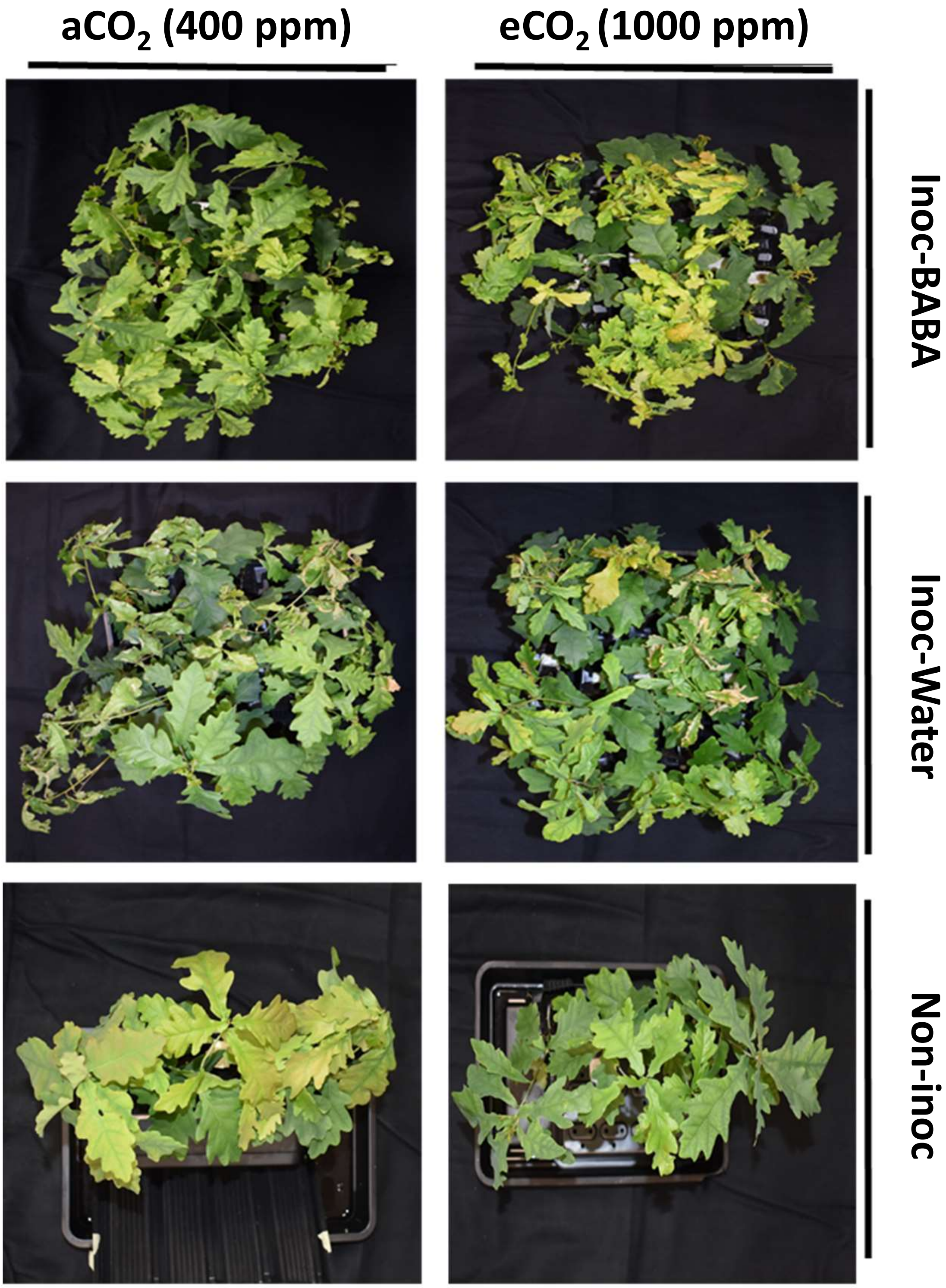
